# The amino acid permease Byc1 is involved in calcineurin-dependent thermos-tolerance in *Cryptococcus neoformans*

**DOI:** 10.1101/2024.02.27.582329

**Authors:** Kyung-Tae Lee, Madeline Giner, Daniel Bahn, Courtney P. Cornitius, Soo Chan Lee

## Abstract

Cryptococcosis is one of the most serious infectious fungal diseases. It is known to infect the human respiratory tract and central nervous system and cause meningitis and encephalitis, especially in patients with AIDS. In the sub-Saharan area, cryptococcosis causes more mortality than tuberculosis. *Cryptococcus neoformans* is the etiological agent. There has been a growing number of reports of antifungal resistance in *C. neoformans*, and this fungus is also primarily resistant to the newest antifungal drug class, the echinocandins. Despite advances in modern medicine, the morbidity and mortality of *C. neoformans* infections remain unacceptably high. In *C. neoformans*, calcineurin, a serine/threonine phosphatase, is required for growth at the human host temperature, and calcineurin mutants are not virulent in murine models of cryptococcal meningitis. Therefore, calcineurin has long been considered an antifungal target. However, calcineurin is highly conserved in fungi and humans, and thus direct pharmacological inhibition of calcineurin can risk lowering host immunity as the human calcineurin is required for T-cell responses. A previous study demonstrated that there is a link between calcineurin, an amino acid permease, and protein kinase A’s in Mucorales. In this study, we further reveal the existence of a novel link between calcineurin and an amino acid permease, Byc1, for a thermo-tolerance phenotype in this fungus. We found that calcineurin negatively regulates the expression of the *BYC1* gene and that *byc1*Δ mutants are partially resistant to FK506. Moreover, we revealed that overexpression of the *BYC1* gene recapitulates the calcineurin mutants’ thermos-sensitive phenotypes, including FK506 sensitivity and lowered virulence in a murine infection model. We also found that protein kinase A (Pka1) mutants are also partially resistant to FK506. These results support the link between calcineurin, Byc1, and Pka1 in *C. neoformans* and will serve as a platform for developing novel antifungal drugs against the calcineurin pathway without directly targeting the phosphatase.

## Introduction

In recent decades, life-threatening fungal infections have become increasingly prevalent due to the growing number of susceptible immunocompromised cohorts. Cryptococcosis is one of the most serious infectious fungal diseases and is known to target the human respiratory tract and central nervous system, especially in patients with AIDS. In the sub-Saharan area, cryptococcosis causes more mortality than tuberculosis (20). *Cryptococcus neoformans* is the etiological agent. The disease can be treated with an azole such as fluconazole, but cryptococcal meningitis may require the use of polyenes such as amphotericin B (21). Clinically, the combination of amphotericin B with flucytosine has resulted in better outcomes than treatment with amphotericin B alone (5, 11). Regardless, there has been a growing number of reports of antifungal resistance in *C. neoformans* (1, 7, 15, 23), and the fungus is also primarily resistant to the newest antifungal drug class, the echinocandins. Despite advances in modern medicine, the morbidity and mortality of *C. neoformans* infections remain unacceptably high, with mortality rates of up to 20% in infected AIDS patients. Thus, there is no doubt that cryptococcosis poses a significant threat to the health care system and that the development of alternative treatment strategies is urgently needed.

Calcineurin is a serine/threonine phosphatase and has long been considered an antifungal target. In *C. neoformans* (26), calcineurin is required for growth at the human host temperature and the calcineurin inhibitor FK506 is toxic to the fungus at 37°C (17, 18). As a consequence, calcineurin mutants are not virulent in murine models of cryptococcal meningitis (18). There are two well-known calcineurin inhibitors: FK506 and cyclosporine A (CsA). FKBP12 is an immunophilin family protein with a *cis-trans* peptide prolyl isomerase activity that serves as a cellular receptor for the calcineurin inhibitor FK506 (22). When bound to FK506, FKBP12 binds to the interface between the calcineurin catalytic A and regulatory B subunits, inhibiting phosphatase activity by occluding substrate access to the active site (14). Another immunophilin, cyclophilin A (Cyp), also serves as a cellular receptor for cyclosporine A (CsA). CsA-Cyp inhibits calcineurin by blocking the access of the substrates to the active site. However, calcineurin is highly conserved in fungi and humans, and thus direct pharmacological inhibition of calcineurin can risk lowering host immunity as the human calcineurin is required for T-cell responses. Indeed, FK506 and CsA are clinically used to suppress host immunity to avoid rejection in solid-organ transplant recipients. Therefore, understanding downstream targets, such as the downstream factors of calcineurin that play a role in growth at 37°C, and their role in virulence are necessary.

A study revealed for the first time that an amino acid permease is a downstream factor in the calcineurin pathway in the model Mucorales, *Mucor circinelloides* (hereafter *Mucor*). In *Mucor*, the *bycA* gene encoding an amino acid permease is a downstream factor of the calcineurin involved in invasive hyphal growth (25). In brief, calcineurin negatively regulates the expression of the *bycA* gene, and the elevation of *McbycA* gene expression results in a lack of hyphal growth that exhibits yeast-locked growth in *Mucor*. The study also found that the *McbycA* gene links calcineurin and protein kinase A (PKA), in which FK506-sensitive or calcineurin-defective phenotypes are expressed via elevated PKA activity.

In this study, we show that the link between calcineurin, an amino acid permease, and Pka1 also functions in thermos-tolerance in *C. neoformans*. In brief, calcineurin negatively regulates the expression of *byc1* that encods an amino acid permease, and *byc1*Δ mutants are partially resistant to FK506. In addition, *pka1*Δ mutants are also partially resistant to FK506. This study also reveals a potential transcription factor(s) linking calcineurin and *byc1*. More importantly, overexpression of *byc1* resulted in the inability of the fungus to grow at 37°C, and this is a recapitulation of calcineurin inhibitions without direct compromising calcineurin functions. These findings shed light on novel antifungal drug developments targeting Byc1.

## Results and discussion

### Conserved link between calcineurin and Byc1 in *Cryptococcus neoformans*

In *Mucor*, calcineurin governs hyphal growth, in which the phosphatase negatively regulates the expression of the amino acid permease gene *McbycA*. Thus, when calcineurin is inhibited, the expression of *McbycA* is elevated and protein kinase As’ (PKAs’) activity increases and hyphal growth is suppressed with yeast-locked morphology (25). We hypothesized that this link is also conserved in *C. neoformans*. This led us to identify a *C. neoformans* homolog of the *Mucor* BycA. Although the *C. neoformans* (the H99 strain) genome encodes eight amino acid permeases (16), none of them actually shares sequence similarity (data not shown); the blast search against the *C. neoformans* genome with the *Mucor* BycA comes up with no homologs. To identify a potential McBycA-equivalent amino acid permease in *C. neoformans*, we focused on the expression pattern, expecting that the gene expression of the candidate(s) would be negatively regulated by calcineurin.

Chow et al. revealed groups of genes that are differentially regulated in the presence and absence of calcineurin in a Crz1-independent way (2). They found that the expression of a gene encoding an amino acid permease (CNAG_01118 or Aap3) met the criteria similar to *bycA* in *Mucor* (25) (designated as *BYC1* in this study). Our RT-PCR analysis further verified that the expression of *BYC1* was elevated in the wild-type (WT) H99 cells when incubated for 24 hours at 37°C in the presence of FK506 compared to the non-FK506 condition (Fig 1). Three independent experiments with two different pairs of primers for each gene were conducted with similar results, and each experiment was repeated three times.

**Figure 1.**
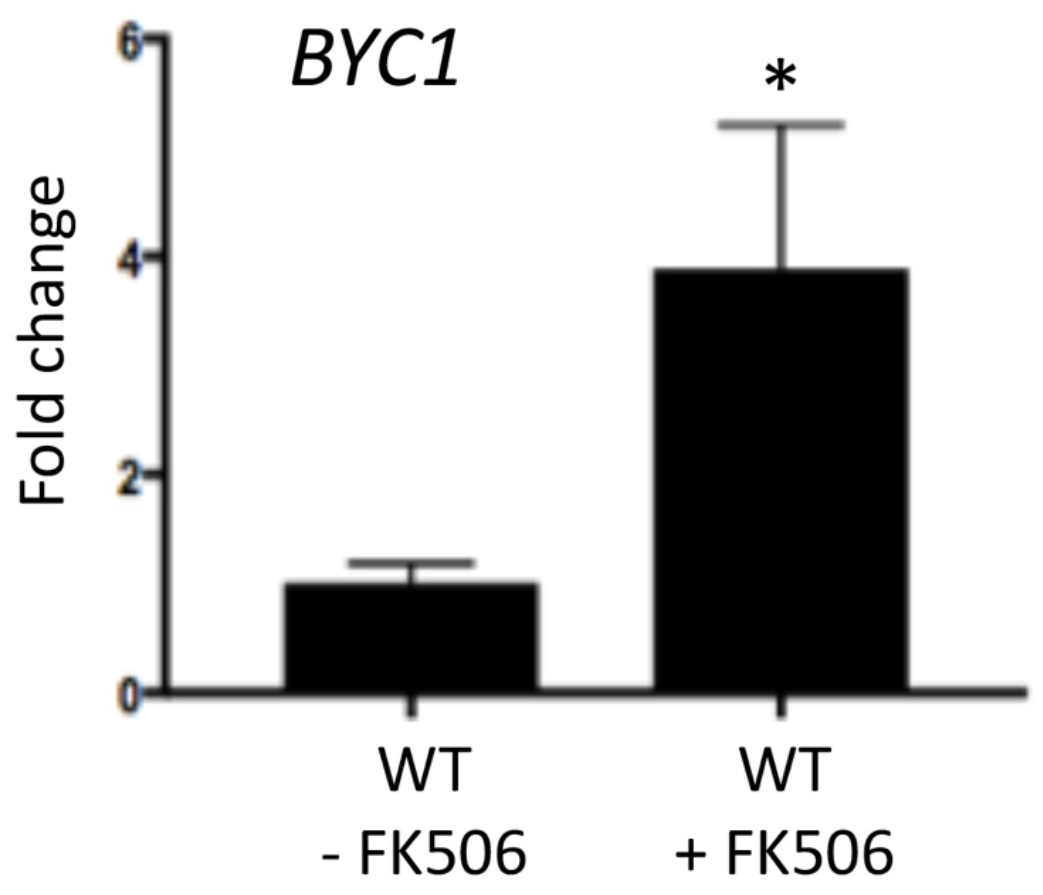
Expression of *BYC1* in the presence of FK506. Inhibition of calcineurin by FK506 treatment results in an elevation of the expression of *BYC1*, indicating that calcineurin negatively regulates the expression of *BYC1*.

The *BYC1* gene was replaced with a neomycin marker, and the deletion of the genes was confirmed by PCR and Southern blotting (data not shown). *byc1*Δ mutants were then grown in YPD media in the presence or absence of FK506 (Fig 2). Interestingly, the *byc1*Δ mutants exhibited partial resistance to the calcineurin inhibitor FK506, whereas WT was sensitive to FK506 at 37°C. When the *BYC1* gene was deleted in the calcineurin catalytic A subunit (*cna1*Δ) or regulatory B subunit (*cnb1*Δ) mutant background, the *cna1*Δ *byc1*Δ and *cnb1*Δ *byc1*Δ exhibited partial thermo-tolerance at 37°C, whereas the calcineurin catalytic A or regulatory B subunit mutants (6, 19)were thermo-sensitive at 37°C (Fig 2). It appears that, in *C. neoformans*, calcineurin regulates thermos-tolerance via Crz1-dependent (2) and Crz1-independent Byc1 pathways.

**Figure 2.**
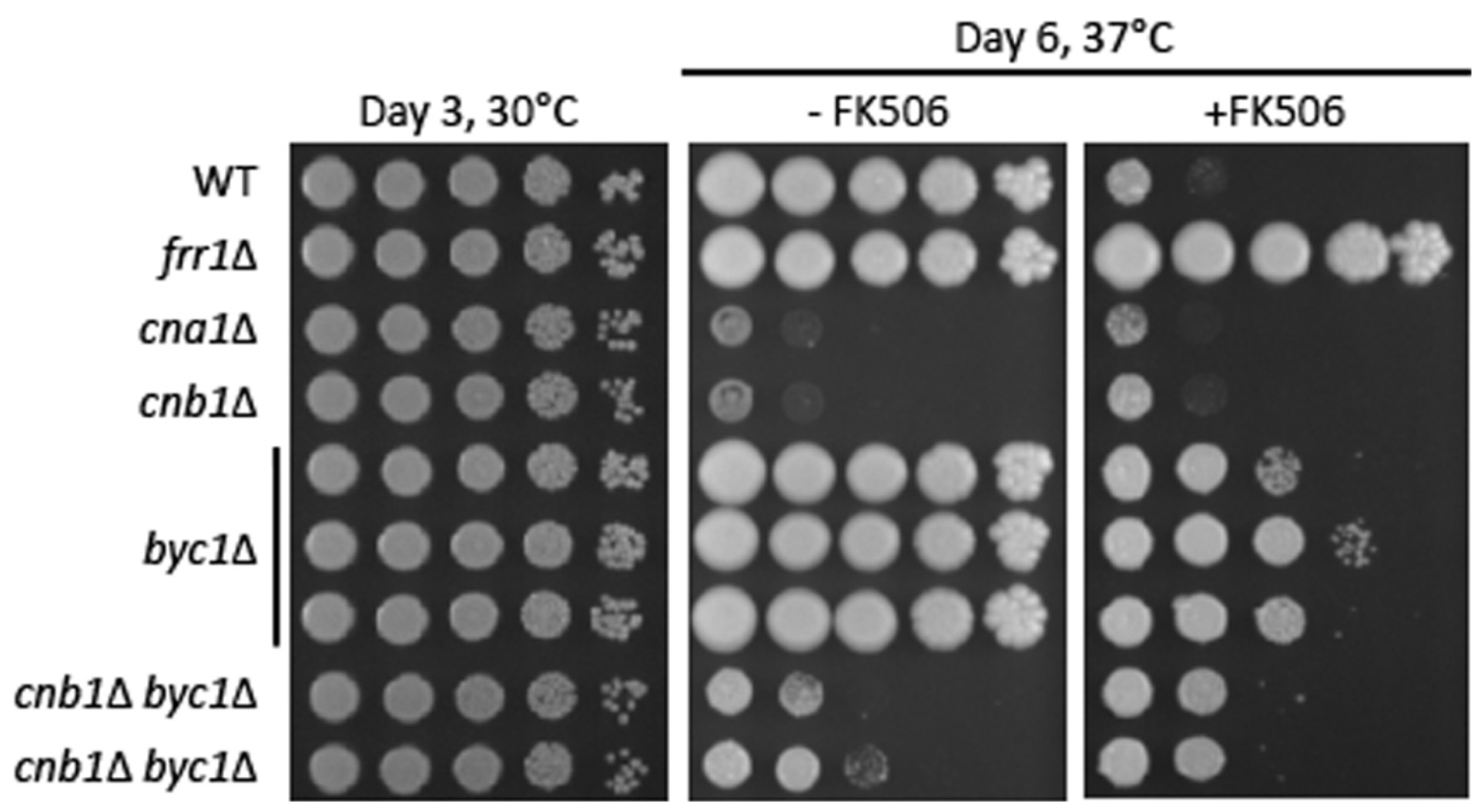
The *BYC1* gene is involved in thermo-tolerance in *C. neoformans. byc1*Δ, *cnb1*Δ *byc1*Δ, and *cna1*Δ *byc1*Δ mutants were spotted onto YPD containing or not containing FK506 and incubated for 6 days at 37°C. Three independently obtained *byc1* mutants exhibited partial resistance to FK506 compared to WT, and *cnb1*Δ *byc1*Δ and *cna1*Δ *byc1*Δ mutants also exhibited partial thermo-tolerance compared to *cna1*Δ or *cnb1*Δ. The *FRR1* gene encodes a FKBP12 that is a cellular receptor for FK506, and lack of this gene therefore results in resistance to FK506.

### Deletion of *BYC1* results in bypass of calcineurin mutants in sexual development

In *C. neoformans*, calcineurin is also known to function in sexual development. *cna1*Δ and *cnb1*Δ mutants exhibit substantial defect of filamentation (3, 6, 19). We found that *BYC1* is also involved in this function of calcineurin (Fig 3). The *BYC1* gene was deleted in addition to *cna1*Δ or *cnb1*Δ in an α mating type cell. When *cna1*Δ or *cnb1*Δ was co-cultured with a WT **a** cell, the calcineurin mutants did not undergo filamentation observed in **a**WT x αWT. However, when the *BYC1* gene was deleted on top of the calcineurin genes and the double mutants *cna1*Δ *byc1*Δ or *cnb1*Δ *byc1*Δ were co-cultured with **a**WT, higher filamentation was observed compared to the single calcineurin mutants. These results also indicate that *C. neoformans* Byc1 is also involved in sexual development.

**Figure 3.**
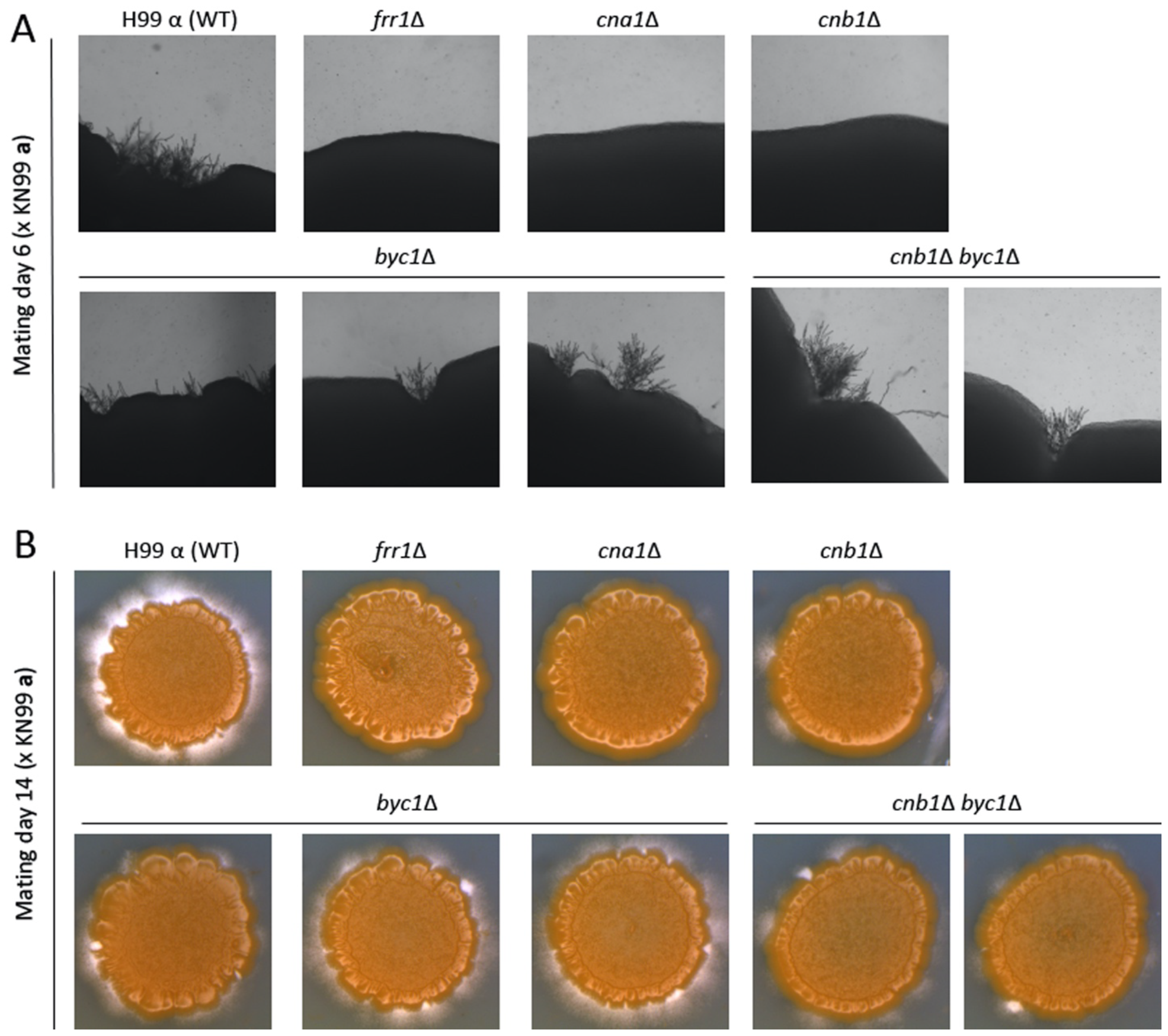
The *BYC1* gene is involved in sexual development. *byc1*Δ, *cnb1*Δ *byc1*Δ, and *cna1*Δ *byc1*Δ mutants were co-cultured with KN99**a** strain. The calcineurin mutants (frr1Δ, *cnb1*Δ, *cna1*Δ) did not form robust filamentations when co-cultured with KN99**a**, unlike the mating between wild type and KN99**a**. However, *byc1*Δ mutations in the background of *cna1*Δ or *cnb1*Δ did indeed bypass the filamentation defects.

### Overexpression of *BYC1* in WT phenocopies of calcineurin mutants in virulence

Based on these findings, we further hypothesized that overexpression of *BYC1* would result in elevated thermo-sensitivity at 37°C (Fig 4). To test this, we generated a construct with the *BYC1* gene under a histone H3 promoter (*P*_*H3*_::*BYC1::NAT*) (note: *NAT* is a nourseothricin-resistant cassette), which would constitutively overexpress the downstream gene. The construct was synthesized (GenScript), including the *BYC1*’s own terminator at the 3’-end. The linearized construct was introduced into the genome of the WT *C. neoformans* H99 strain by electroporation (13). The overexpression of *BYC1* in three independent *P*_*H3*_::*BYC1* mutants was confirmed via qRT-PCR (Fig 4B) and the strains exhibited growth defect at 37°C, similar to calcineurin mutants (Fig 3A). More importantly, *P*_*H3*_::*BYC1 -1* is significantly lower in virulence (Fig 4C) compared to WT, similar to calcineurin mutants (19). Balb/c mice (18 to 20g) were used as a host. Mice were anaesthetized with isoflurane (2%) and infected via intratracheal route (1 X 10^6^ cells) suspended in 50 μl of sterile PBS. This result strongly supports the link between calcineurin and *BYC1* regarding thermo-tolerance and virulence.

**Figure 4.**
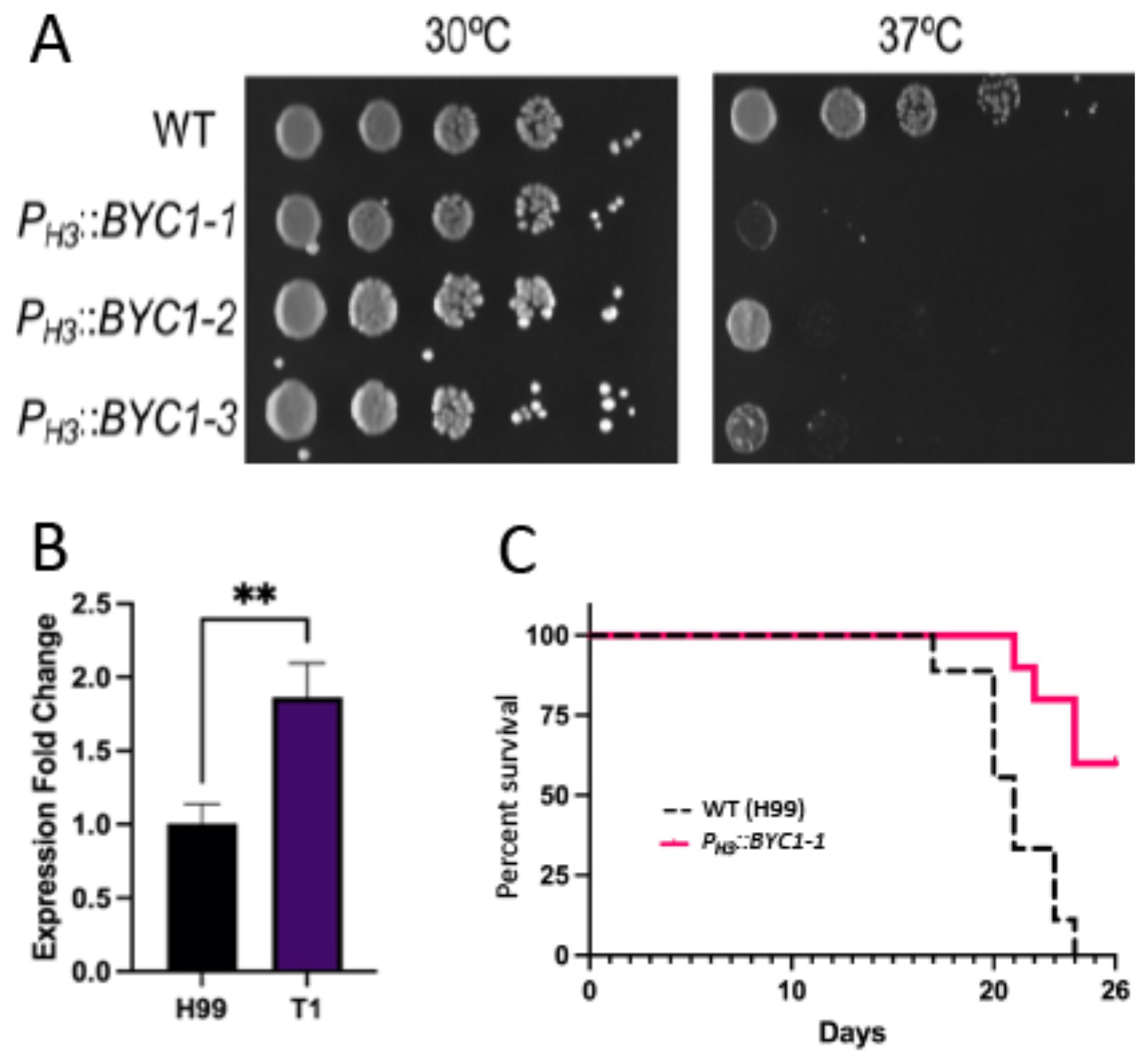
Overexpression of *BYC1* results in growth defect at 37°C (A) and lower virulence (C). The *BYC1* expression is elevated in T1 transformant, indicating that *P*_*H3*_*::BYC1* does indeed cause the overexpression of *BYC1*. Three independently derived transformants with *P*_*H3*_::*BYC1* exhibit similar growth defects at 37°C (A). Overexpression of *BYC1* results in significantly lower virulence (p=0.0007).

### Transcription factors involved in FK506 resistance and the expression of *BYC1*

It is highly important to understand the expression mechanism of *BYC1* in *C. neoformans* as the gene can be an alternative target for developing antifungal drugs that are related to calcineurin but do not directly target the phosphatase. It remains to be determined how calcineurin regulates the expression of the *BYC1* gene in *C. neoformans* as the expression of *BYC1* is independent of the known transcription factor Crz1 (2). Thus, to identify key transcription factor(s) in the link between calcineurin and Byc1, we screened a transcription factor knockout library (10). We found that a transcription factor mutant (CNAG_00018) exhibited partial resistance to FK506 at 37°C compared to WT (Fig 5A). We also verified that the expression of *BYC1* is significantly lowered in the Δ*CNAG_00018* mutant (Fig 5B). Two independently derived mutants (10) exhibited similar phenotypes.

**Figure 5.**
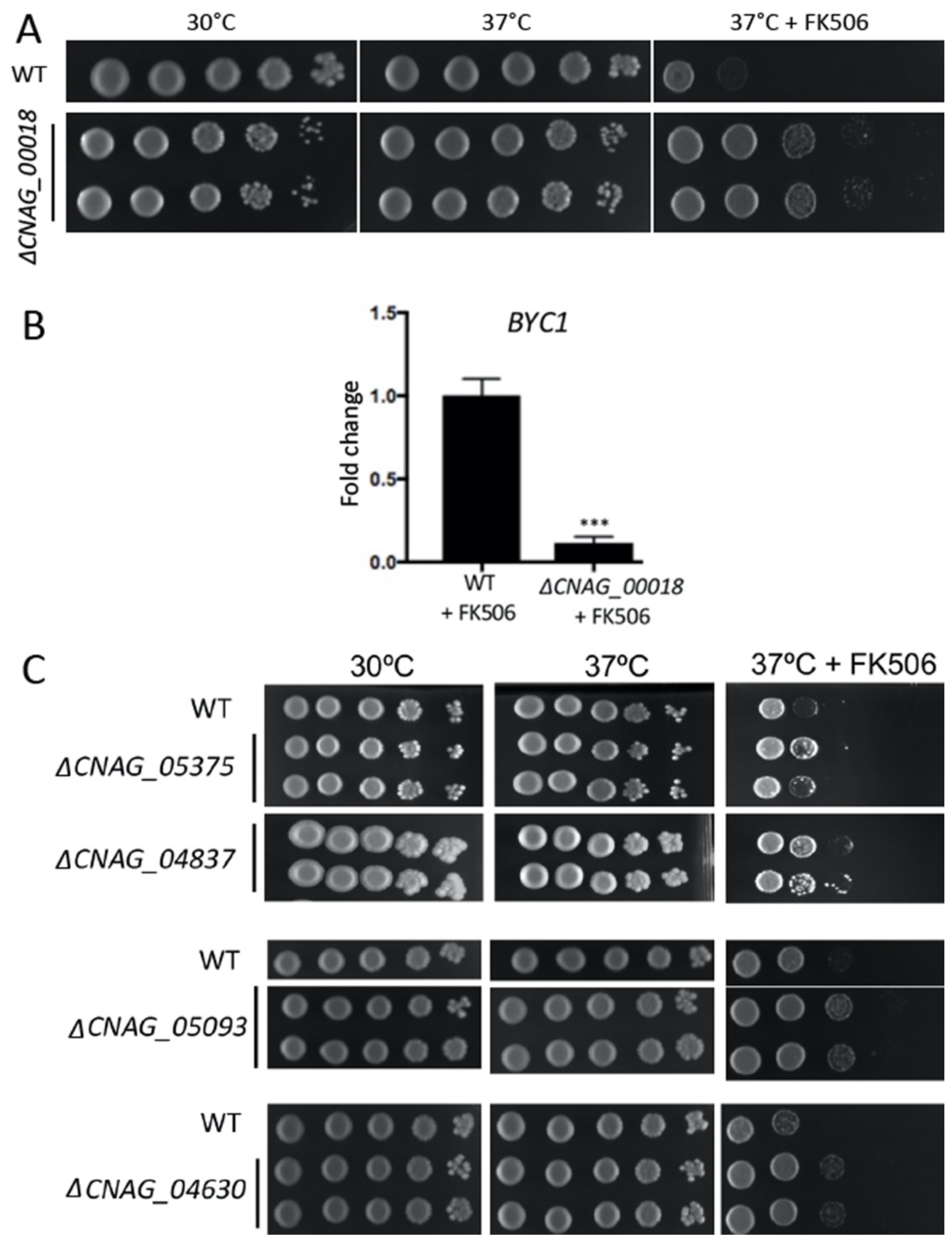
Deletion of the gene encoding a transcription factor (CNAG_00018) resulted in partial thermo-tolerance at 37° C in the presence of FK506 with two independently derived mutants (A). In the mutants, the *BYC1* expression was lowered and was not induced by FK506 (B). Deletion of the gene encoding a transcription factor (CNAG_00018) resulted in thermo-tolerance at 37°C in the presence of FK506 with two independently derived mutants (C).

**Figure 6.**
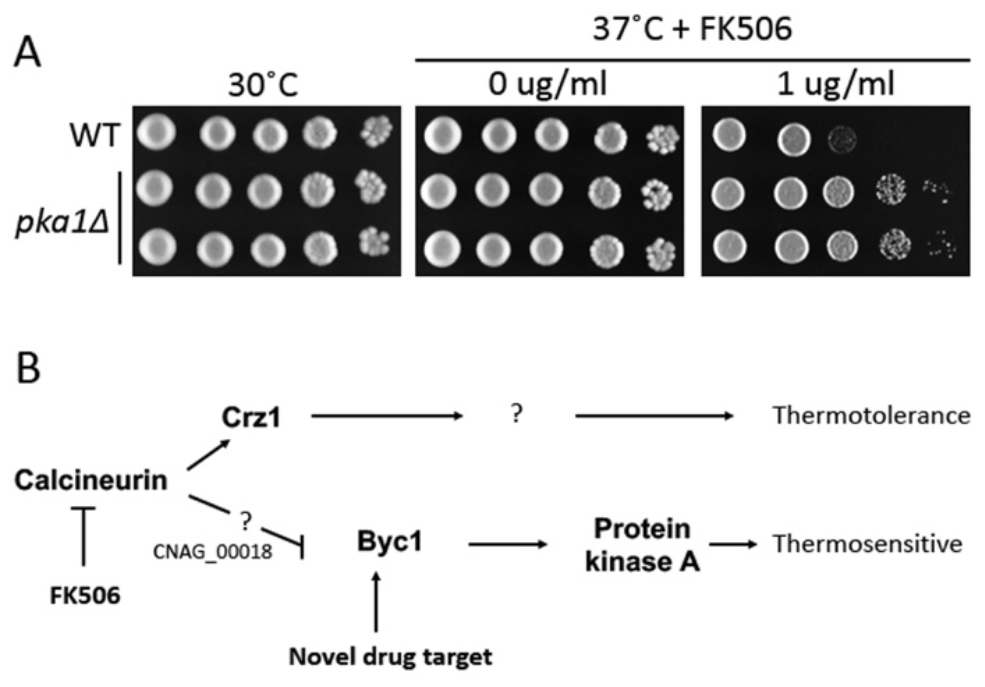
Protein kinase A (PKA) is involved in thermo-sensitivity when calcineurin is inhibited. Two independently derived pka1 mutants exhibited resistance to FK506 at 37°C (A). (B) depicts the overall scheme of calcineurin, Byc1, and PKA in thermos-tolerance. It is suggested that targeting Byc1 expression can be used to develop novel antifungal drugs without directly inhibiting calcineurin.

This suggests that CNAG_00018 can be a linking transcription factor between calcineurin and *BYC1*. For example, calcineurin dephosphorylates CNAG_00018, in which the transcription factor is in inactive status, and in turn the expression of *BYC1* is not turned on. On the other hand, calcineurin in inhibited, active phosphorylated CNAG_00018 can turn on the *BYC1* gene, exhibiting a heat-sensitive phenotype. Alternatively, CNAG_00018 may serve as a transcription factor for a negative regulator of the BYC1 gene. For example, activating CNAG_00018 by dephosphorylation by calcineurin can trigger the expression of the *BYC1* repressor. When calcineurin is inhibited, CNAG_00018 becomes inactive and the repressor is not expressed. Therefore, *BYC1* is highly expressed. More experiments will be needed to unveil the exact mechanism. There are more transcription factors potentially involved in this mechanism, including CNAG_05375, CNAG_04837, CNAG_05093, and CNAG_04630. These mutants are also partially resistant to FK506. A transcription factors–*BYC1* promoter binding assay will verify whether they are a direct transcription regulator of the *BYC1* gene. As *BYC1* expression regulation is a fungal-specific calcineurin pathway inhibition target for drug development, it is essential to understand how these transcription factors are linking calcineurin and the *BYC1* gene at the molecular level.

### Mutations in the protein kinase A gene confer resistance to a calcineurin inhibitor

Our previous study (25) revealed that calcineurin is antagonistic to the PKA activity in *Mucor* and *C. neoformans*. In *Mucor*, lack of calcineurin results in cAMP independent activation of PKAs (25). In addition, our previous study found that PKA activity in *C. neoformans* is also antagonistic to calcineurin (12). This prompted us to test whether *C. neoformans pka* mutants are resistant to calcineurin inhibitor FK506. Interestingly, pka1 mutants are partially resistant to FK506, which is a phenocopy of *byc1* deletions and CNAG_00018 deletions. Thus, this result demonstrates that the link between calcineurin and PKA is extant in *C. neoformans*.

However, *PKA2*, another gene-encoding PKA catalytic subunit, appeared not to be involved in the link with calcineurin as *pka2Δ* mutants are as sensitive to FK506 as WT. In *Candida albicans*, the two isoforms of PKA catalytic subunits Tpk1 and Tpk2 play different roles in stress response and carbohydrate metabolism (8). In humans, some PKA catalytic subunit isoforms localize on membranes and become active, representing distinct roles of each isoform (9). It is also possible in *C. neoformans* that two isoforms of PKA catalytic subunits (Pka’s) have a separate function.

In *S. cerevisiae*, the general amino acid permease Gap1 is known to function as a nutrition sensor and to activate PKA in a cAMP-independent manner (4). The amino acid permease has dual functions, including signaling to PKA and amino acid transportation. How amino acid permease activates PKA is therefore of interest. It has been further suggested by a previous study that the C-terminus of Gap1 functions in PKA activation and has a consensus motif (I/L/φ)xxxxx(I/L/φ)xxTKxxxxxxφxxφ (where φ is a hydrophobic amino acid), shared with a protein kinase A regulatory subunit (Bcy1) (24). It is therefore hypothesized that the C-terminus motif of Byc1 in *C. neoformans* activates Pka1 by interfering the inhibitory function of the protein kinase A regulatory (PKR) subunit, which needs to be addressed for a complete understanding of the calcineurin-Byc1-Pka link.

## Acknowledgments

We thank Minjae Lee, Seungheon Lee, Sandeep Vellanki, Alex Ko, and Alex Garcia for technical support. We are indebted to J.H. for providing calcineurin mutant strains. S.C.L holds a Voelcker Fund Young Investigator Award from the Max and Minnie Tomerlin Voelcker Fund.

## Notes

### Competing Interest Statement

The authors have declared no competing interest.

